# Mitigating Data Scarcity in Protein Binding Prediction Using Meta-Learning

**DOI:** 10.1101/519413

**Authors:** Yunan Luo, Jianzhu Ma, Xiaoming Zhao, Yufeng Su, Yang Liu, Trey Ideker, Jian Peng

**Affiliations:** Department of Computer Science, University of Illinois at Urbana-Champaign; School of Medicine, University of California San Diego; School of Electronic Information and Electrical Engineering, Shanghai Jiao Tong University

## Abstract

A plethora of biological functions are performed through various types of protein-peptide binding. Prime examples include the protein kinase phosphorylation on peptide substrates and the binding of major histocompatibility complex to neoantigens in the immune system. Understanding the specificity of protein-peptide interactions is critical for unraveling the architectures of functional pathways and the mechanisms of cellular processes in human cells. Despite mass-spectrometric techniques were developed for the identification of protein-peptide interactions, our understanding of the preferences of proteins on their binding peptides is still rudimentary. As a complementary direction, a line of computational prediction methods has been recently proposed to predict protein-peptide bindings which efficiently provide rich functional annotations on a large scale. To achieve a high prediction accuracy, these computational methods require a sufficient amount of data to build the prediction model. However, the number of experimentally verified protein-peptide bindings is often limited in real cases. For example, a majority of protein kinases have very few experimentally verified phosphorylation sites (e.g., less than 30 sites) in existing databases. These methods are thus limited to building accurate prediction models for only well-characterized proteins with a large volume of known binding peptides and cannot be extended to predict new binding peptides for less-studied proteins. In this paper, we introduce a generic framework to address this issue of data scarcity in protein binding prediction. We demonstrate the applicability of our framework in predicting kinase-specific phosphorylation sites. Our method uses an effective training strategy to build a prediction model with robust transferability. The model is able to predict the phosphorylation sites of a less-studied kinase, even if there is only a small number of phosphorylation sites known for this kinase. To achieve this, we train the model via a meta-learning phase followed by a few-shot learning phase. We demonstrate our framework has better transferability than state-of-the-art methods and is effective in utilizing limited data to accurately predict phosphorylation sites for less-characterized kinases. The implementation of our framework is available at https://github.com/luoyunan/MetaKinase.

## 1 Introduction

Proteins interact with signaling molecules called ligands to induce a change in biological activities. The bonds at the binding site are noncovalent and transient to easily regulate the activation of certain biological functions. Characteristics of binding specificity measure the types of ligands, such as peptides or DNA/RNA, that a protein will bind to. As a prime example, protein phosphorylation is the major molecular mechanism involved in a variety of fundamental cellular processes such as proliferation, differentiation, and apoptosis [1]. Phosphorylation requires the physical interaction between a protein kinase and a short peptide, in which the protein kinase adds a phosphoryl group to the phosphorylated residue in a peptide. The human genome encodes more than 500 unique kinases which can be organized in a hierarchy of groups, families, and subfamilies [2], and many of them have evolved to have highly divergent specificities [3]. Despite the recent advances in high-throughput mass-spec techniques, it is still costly and challenging to experimentally characterize the specificity of kinase families due to the high specificity diversity, making it a pressing need to develop computational methods that are capable of capturing kinase specificities. Such challenges exist in many protein-ligand binding problems, such as for other signaling proteins and MHC proteins.

Previous computational models have mainly followed the “one-family-one-model” paradigm to address the challenge of characterizing divergent specificities of different kinase families. That is, for a given kinase family, a separate machine learning model was built to capture the unique pattern embedded in the substrate sequences of this family. Many successful applications belong to this family-specific category, such as NetPhosK (Neural Networks) [4], KinasePhos (Hidden Markov Models) [5], PhosPhoPICK (Bayesian Networks) [6], PhosphoPredict (Random Forests) [7], MusiteDeep (Convolutional Neural Network) [8], and many others [9–13]. An important prerequisite of these family-specific methods is the availability of a sufficient amount of high-quality data for model training. However, this is often not the case: a majority of kinase families have less than 30 experimentally verified phosphorylation sites. As a result, previous studies have to choose to build family-specific models for well-characterized kinase families (typically <10 families) with abundant training data [7,8,14]. For those less-studied kinase families, using the family-specific approach fails to provide satisfactory performance due to the lack of sufficient training data. The data sparsity thus raises a challenge for the prediction of protein-peptide interactions, as data-driven approaches highly rely on a large quantity of data to achieve accurate predictions.

To alleviate the data scarcity problem in predicting phosphorylation sites of less-characterized kinases, recent works have proposed to use a “pan-family” approach to build the prediction model [15, 16]. The idea of this type of approaches is that there exist universal patterns of binding affinity shared by all kinase families and they could be potentially transferred to benefit the prediction of less-studied kinase families by integrating all the other kinase families. Specifically, besides peptide sequences, the pan-family approach also needs to consider kinase sequences as an additional input to build the universal model for all kinase families. By leveraging the data from all families, the pan-family model is able to produce accurate predictions for kinases with limited measured data. The pan-family approach has also been applied in modeling other types of protein-peptide interactions, such as the prediction of binding affinity of major histocompatibility complex (MHC) to neoantigens [17]. While having been demonstrated to have the superior transferability and be able to improve the prediction performance of less-characterized families [15], pan-family approaches still suffer a major limitation: the single model is generally hard to express the specificities of all protein families given limited model complexity. Therefore, it is reasonable to expect that pan-family approaches may achieve a worse prediction accuracy than a family-specific model for a particular family with a relatively large number of training data. How to capture both transferability across families and family specificity simultaneously has becomes a new challenge in the prediction of protein-peptide interactions. To our best knowledge, no previous work has taken the initiative to tackle this challenge.

As discussed above, the “one-family-one-model” design is able to accurately characterize specificities of individual families with a large volume of training data but cannot be generalized to less-studied families. Pan-family approaches can integrate available data of all families but may lose fine-grained family-specific patterns by merging different families together. To address the problems of both models, in this work, we propose a new two phases *meta-learning* framework, named MetaKinase, for the prediction of kinase phosphorylation sites. In phase one, using multiple training kinase families, we train a model which can generate more adaptable representations which are broadly suitable for every kinase family (called meta-learning). In phase two, using a few (e.g., < 10) known phosphorylation sites from a new target kinase family, we fine-tune the model on this target family to capture its specificity. With the general patterns captured in phase one, the adaption to the target family in phase two is very sample-efficient: we can tweak the model by only using a few data points to make it family-specific and accurately predict the specificity of the target family (called few-shot learning). With its transferability and fast adaptability, our framework can thus be applied to mitigate the data scarcity issue in characterizing specificities of less-studied kinases. Even with only a few known phosphorylation sites, the model is still able to accurately characterize the specificity of the target kinase family.

Apart from previous works, including both family-specific and pan-family approaches, that primarily focused on the designing of model architecture itself, our work illustrates how to address the data scarcity challenge in protein-peptide interaction prediction in a new direction, i.e., the way of training the model. In addition, our framework is model-agnostic: in principle, the framework can take any model as its the base predictor, as long as the model can be optimized using gradient descent (e.g., neural network). This includes a wide range of models, including existing predictors of kinase-specific phosphorylation sites such as DeepSignal [15] and MusiteDeep [8]. In this work, we develop a simple convolutional neural network (CNN) as the base predictor. We demonstrate that MetaKinase is capable of accurately predict the phosphorylation sites of a new kinase using only a few data samples of this kinase. Experiment results show that our framework is effective and significantly outperforms existing approaches for phosphorylation site prediction when training data is limited. We also show that our framework is sample-efficient and can quickly capture the specificity of the test kinase.

## 2 Data

### 2.1 Phosphorylation sites

We constructed a dataset of 15-mer peptides to assess the performance of our framework in comparison with other methods. Peptides in the dataset all contain the phosphorylation sites (either Serine (S), Threonine (T) or Tyrosine (Y)) centered in the middle position of the 15 amino acids. Although longer peptides may improve the prediction performance as reported in [8], length 15 was widely adopted in previous works [11, 15, 18] so here we just follow this setting without further optimization. These peptides were pulled from public databases UniProt [19] and Phospho.ELM [20], following the procedure described in [14]. We downloaded reviewed proteins that contain at least one human phosphorylation sites from UniProt and removed phosphorylation sites without annotated up-stream kinase information. In addition to the UniProt database, phosphorylation sites from the Phospho.ELM database were also collected. We removed the redundant entries in both of the two databases and eventually obtained 2, 479 S sites, 657 T sites and 1, 202 Y sites that covered 46 kinases of 25 kinase families.

### 2.2 Kinase sequences

The amino acid sequences of human kinase domains were retrieved from KinBase [21]. These domain sequences were searched using HHblits [22] and re-aligned using HHalign [23] with their default parameter setting. The multiple sequence alignment was then manually refined by removing alignment gaps at both ends. We grouped individual kinases into 25 kinase families based on the hierarchical kinome classification system [21].

### 2.3 Training and test set

To create the training and test set, for each kinase we randomly divided our downloaded proteins containing phosphorylation sites into five subsets, where four subsets were used as positive training samples and the remaining subset was used as positive testing samples. Negative samples were obtained by extracting S/T/Y sites from the same protein sequence in the training set and the test set respectively and removing those sites annotated as phosphorylation sites. Here, we also matched the type phosphorylation residues of a kinase in both the positive samples and negative samples. For example, if a kinase only phosphorylates Y sites, we only extracted background Y sites as negative samples for this kinase, excluding S sites and T sites.

## 3 Data-driven Motivation

To motivate the design of our framework, we first performed exploratory analysis that leads to two important observations on the data availability of current databases of kinase phosphorylation sites and the transferability of existing methods.

First, we observed that majority protein kinases have a small number of known phosphorylation sites (Figure 1a). We extracted all phosphorylation sites from the Phospho.ELM database that had up-stream kinase annotations [20] (Section 2) and counted the number of known phosphorylation sites of each protein kinase. From the 309 kinases that had at least one phosphorylation site, we found that > 78% of protein kinases had < 30 phosphorylation sites and > 50% had < 10 phosphorylation sites. In contrast, previous family-specific methods for phosphorylation sites prediction often required hundreds of phosphorylation sites to build reliable prediction models to prohibit over-fitting. These methods are thus limited by the current data availability and cannot be used for kinase families with few experimentally verified data. As a result, how to utilize the small amount of data of a less-characterized kinase has become a new challenge, and a model that can integrate available data of other kinases but also capture the specificity of target kinase will be helpful in mitigating the data scarcity issue.

**Figure 1:**
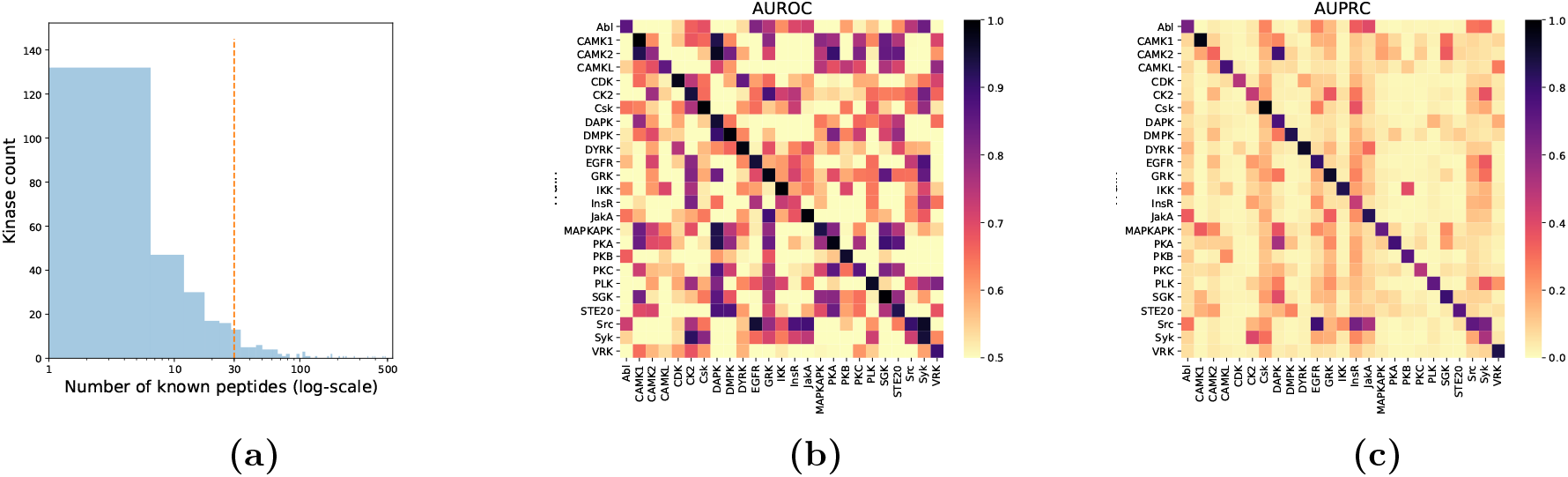
(a) A majority protein kinases have a small amount of known phosphorylation sites. (b)-(c) Family-specific approaches have limited transferability across kinase families.

Second, we observed that family-specific approaches had limited transferability. We bench-marked MusiteDeep, one of the state-of-the-art family-specific, deep learning-based methods of phosphorylation site prediction on the dataset described in Section 2. For each kinase family, we had a non-overlapped training set and test set. We enumerated all possible combinations of the training kinase family and the test kinase family and evaluated the prediction performance using AUROC and AUPRC scores. We found that prediction performance on a test kinase family was maximized if the model was trained using data from the same family. The averaged AUROC when trained on the same kinase family, was significantly higher than the best performance achieved by training on another kinase family (0.93 vs. 0.85; P-value 1.8 × 10^−5^). We observed similar significant gaps in terms of AUPRC (0.73 vs. 0.33; P-value 1.5 × 10^−8^). These results suggested that currently it was difficult to directly apply a model that had been trained on a well-characterized family to predict phosphorylation sites for a less-characterized family. It is important to develop models that have better transferability to improve the prediction for less-characterized kinase families.

## 4 Methods

To enable the prediction of kinase-peptide interactions with only a few samples on the target family, we used a meta-learning – few-shot learning framework to efficiently integrate the information of target family and other kinase families (Fig. 2). We developed a convolutional neural network and trained this network using the meta-learning – few-shot learning strategy, an effective way to better transfer knowledge/features across different kinase families. The trained model was able to quickly adapt to a target kinase family, which possibly was less-characterized, using only a few known interacting peptides of this family. In this section, we will first define the problem setting of meta-learning and few-shot learning and then describe the details of our framework.

**Figure 2:**
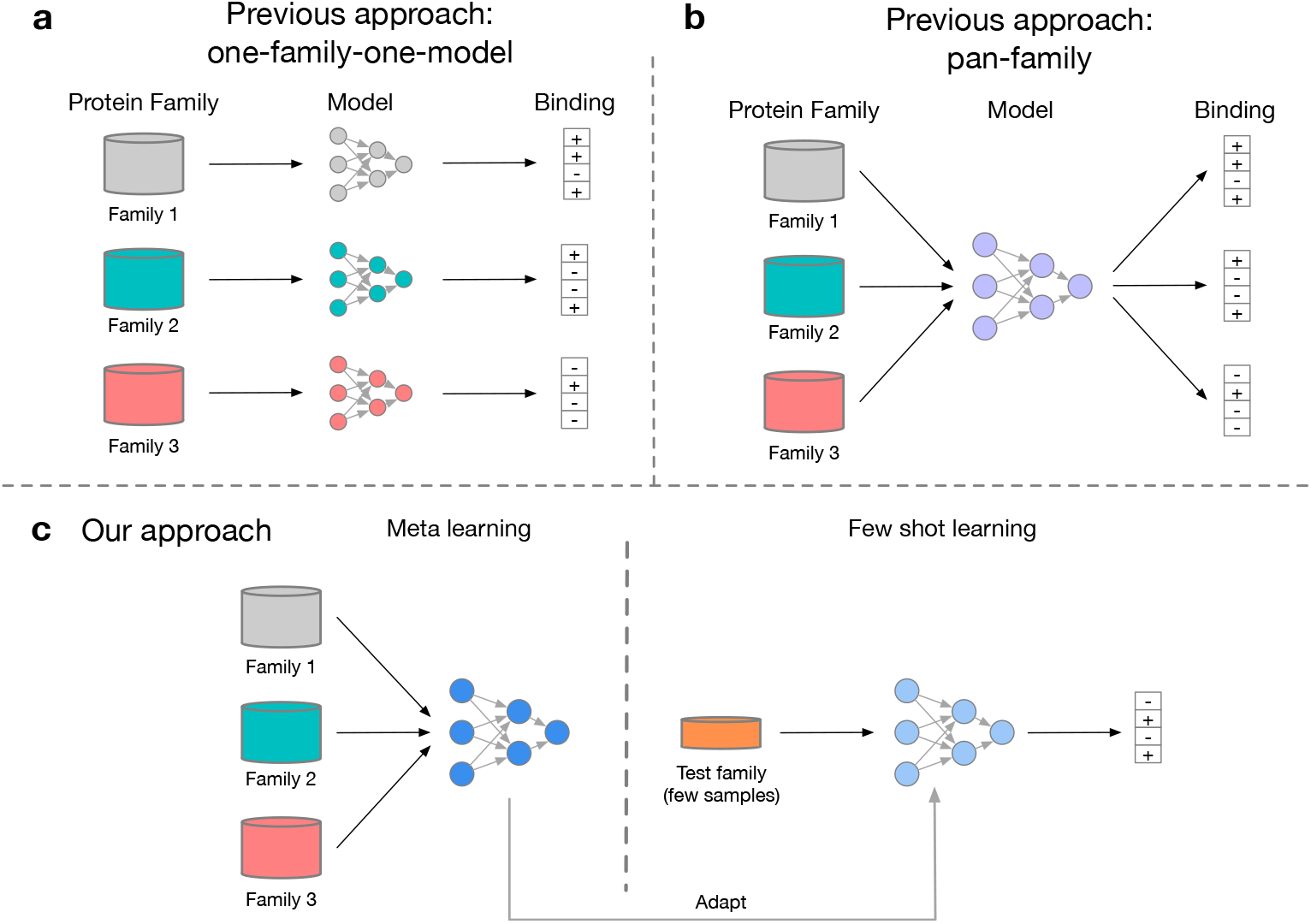
Overview of framework. (a) Previous “one-family-one-model” approach: an individual model is built for a protein family. (b) Previous “pan-family” approach: a single model is built for all protein families. (c) Our approach: the model is first trained on all training families to learn a general representation that is broadly suitable for all families and then fine-tuned using a few data samples of a test family to learn a representation that is specific to the test family.

### 4.1 Overall framework

We are given a set of kinase families 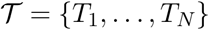 and their corresponding interacting peptides as training data, and *T*\* is the target kinase family that we want our model to characterize its specificity as test data. We also assume we observe a few known interacting peptides of *T*^*^ that can be used as training data. This setting well resembles the scenario of reality: there are several well-characterized kinase families (corresponding to set 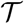) with a sufficient amount of annotated phosphorylation sites verified, which can be used as high-quality training data for data-driven approaches. There are also some completely new or less well-studied kinase families with only a few known phosphorylation sites (e.g., < 10 sites; see Section 3). Our prediction task is to predict new phosphorylation sites for the rest of peptides without measurements based on all the training data from 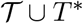.

### 4.2 Meta-learning phase

The key intuition of meta-learning is that there are some representations of data that are more transferable than others. For example, given a set of images, one machine learning model can learn a causal relationship between particular pixels and the labels while another machine learning model can learn the transition-invariant or rotation-invariant properties of the images. The latter is obviously more adaptable to other computer vision applications. Similarly, to encourage a general representation, we employ a meta-learning algorithm [24] to explicitly train a model towards this objective. Technically, the effect of meta-learning is to find model parameters that are sensitive, such that small changes in these parameters can lead to large improvements in characterizing the specificity of a new kinase family.

Formally, let *f_θ_* be the base predictor parameterized by *θ*. For each kinase 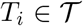, two sets of data points (phosphorylation sites) are sampled from the training data, one set 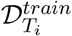 containing *k* samples (e.g., *k* ≤ 10) for training and the other set 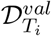 containing *k* samples for validation.

Let 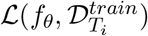 be the training loss (e.g., mean-squared error or cross-entropy error) that the predictor makes on the training samples of kinase *T_i_*. We also have validation loss that defined similarly. In each iteration of the meta-training stage, let us suppose the current model parameter is *θ*. We then sample a batch of kinase families {*T_i_*} from 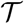, and for each sampled kinase family *T_i_*, we have one or more updates on model parameters which are computed using standard stochastic gradient descent (SGD),

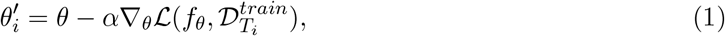

where *α* is the learning rate. After we have an updated parameter 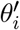 for each test family *T_i_*, we update the original model parameter *θ* by optimizing the performance of 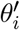 on the validation set 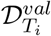 of each kinase family *T_i_*. That is, we minimize the following meta-loss function

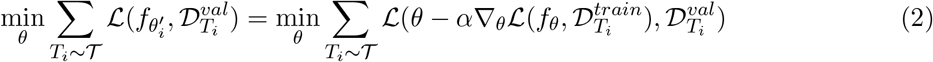

This meta-objective can also be optimized by SGD, in which we have the updating rule

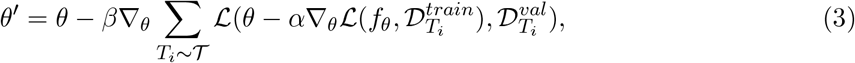

where *β* is the meta-learning rate. After being trained through the above meta-training phase, the model can be applied to predict phosphorylation sites of a new kinase family *T*\*. At the metatesting phase, we assume the target family *T*\* is less-characterized and has only *k* (e.g., *k* ≤ 10) known phosphorylation sites in 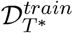. To produce predictions of new phosphorylation site for family *T*\*, we first use the inner-update rule in Eq. (1) to train the model on the *k*-shot samples in 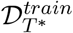, adapting its parameters to the target family and capturing both information shared by multiple families and patterns specific to this family. The model then can be applied to produce predictions for samples in 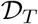.

The meta-learning algorithm we described above requires two passes of back-propagation to compute the Jacobian matrix for optimization, which could be computationally inefficient. In practice, first-order meta-learning is also used for efficiency purpose. We have tested both the vanilla and the first-order version [25] of meta-learning in our in-house experiments and observed similar prediction performances for both methods. We thus chose to use the first-order metalearning algorithm in all presented results in this work as it took a shorter time to train the model.

The kinase-peptide interactions are labeled with a binary variable, indicating whether that kinase phosphorylates the peptide (1) or not (0). We thus use the cross-entropy loss, a natural choice for binary classification, as the loss function in Eq. (2). The dataset of kinase-peptide interactions we collected is highly imbalanced, as for a kinase we often have two orders of magnitude more negative samples than positive samples (Sectioin 2). Therefore, during each iteration of the meta-learning phase, we subsampled negative samples such that the ratio is 1:1 between positive and negative samples. This negative sampling technique was widely used in previous works to address the data imbalance issue [8, 14]. Therefore, the loss of the predictions that a model *f_θ_* makes on the data 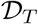 for a kinase family *T* is defined as

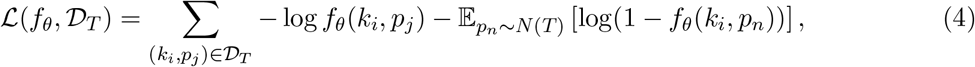

where *k_i_* is a kinase in kinase family *T* and *p_j_* is one of its interacting peptide (i.e., kinase *k_i_* phosphorylates peptide *p_j_*); *p_n_* is a negative peptide of kinse *k_i_*, which is sampled from *N*(*T*), the set of all non-interacting peptides of kinase *k_i_*; *f_θ_*(·) is the model predicted probability that kinase *k_i_* phosphorylates peptide *p_j_*.

To optimize the model, we trained it for 500 meta-iterations. At each iteration, we sampled 40 tasks, with each task being the prediction of *k* random positive samples and *k* random negative samples of a randomly sampled kinase family. Following a previous work [24], we used the Adam optimizer [26] with default learning rate 0.001 to minimize the objective function (Eq. (3)). We set the number of inner-updates to 16 (Eq. (1)). The framework was implemented in PyTorch, and the model was trained on an NVIDIA GTX 1080 GPU.

### 4.3 Few-shot learning phase

In the second phase, we randomly select *k* positive samples and *k* negative samples (*k* varies from one to ten) from the target kinase family as training data to fine-tune the model. The rest positive and negative samples form the testing data. Negative samples are randomly drawn from all S/T/Y sites, excluding annotated phosphorylation sites, from kinase sequences of the target family (Section 2). The model is fine-tuned using the updating rule defined in Eq. (1). This random process is repeated 50 times and the predicting performance is averaged over all the repeats. To gain enough statistical power, we only evaluate the predicting performance on 25 kinase families.

Our framework is able to task a gradient-based model in an arbitrary form as the base predictor, including linear model, Support Vector Machine and various types of Neural Networks. In this work, we choose to use a dual-channel deep neural network as the based predictor (Fig. S1). The input of the predictor is the kinase domain sequence and the peptide sequence centered on the phosphorylation site. We use BLOSUM62 matrix [27] to encode the input sequences. The kinase sequence (~250 amino acids) and the peptide sequence (15 amino acids) are then processed by one of the channels in the neural network, respectively. Each channel is a separate convolutional layer with different lengths of convolution filters, for the purpose of extracting multi-resolution sequence patterns. For the channel of kinase sequence, we used 32 filters of size 3, 5 and 7 amino acids to extract underlying sequence features. A global max pooling operation is performed to select the maximum activation value in each of 32 × 3 = 96 filters and the 96 maximum activation values are concatenated together as the representation of kinase domain sequence. Similarly, the channel of peptide sequence has 32 filters of size 3 and 5 amino acids and the 32 × 2 = 64 maximum activation values are concatenated together as the representation of the peptide sequence. On top of the two channels is a fully-connected layer that concatenates the representations of kinase and peptide sequences to produces the final prediction.

## 5 Results

### 5.1 Experiment settings

Following the few-shot learning setting to evaluate our prediction performance, we performed a leave-one-out experiment on the *N* = 25 kinase families in the collected dataset. In each evaluation, one kinase family was withheld as the test family and the other *N* − 1 families were used as training families. Under the few-shot learning setting, *k* (e.g., *k* ≤ 10) known phosphorylation sites of the test family were also available to the model for training and the remaining samples were used to assess the model performance. This mimics the real scenario that a less-characterized family has very few verified phosphorylation sites available and we want to predict new phosphorylation sites of this family. In the meta-*training* phase, we used the data of the *N* − 1 kinase families to train our model to have a parameter configuration that was broadly suitable for every family. In the meta-*testing* phase, we then fine-tuned the model using the *k* samples of the test family to adapt the model parameters to be specific to the test family. To facilitate the model training in the meta-training and fine-tuning stages, we combined the positive samples with the same number of randomly sampled negative samples (i.e. background peptides centered on non-phosphorylation S/T/Y sites).

### 5.2 Baseline methods

We compared our framework against the following prediction approaches:

- **Pan-family model**: The pan-family approach builds one model for all kinase families. To ensure a fair comparison, we used the same base predictor in our framework for the pan-family approach. The only difference between our framework and the pan-family approach is the training strategy: our framework first trains the model on the *N* − 1 training families through a meta-learning phase and then fine-tunes the model using *k*-shot samples of the target family in the few-shot learning stage, while the pan-family approach trains the model using the *N* − 1 training families as well as the *k* samples from target family together, following the traditional supervised training procedure.
- **K-nearest neighbor (KNN)**: The KNN model first collects all the peptides of *N* − 1 training families together with the *k*-shot samples of the target family. The test peptide is then compared to the *K* most similar training peptides and the prediction follows the majority voting. The peptides were represented using one-hot encoding and the Euclidean distance was used as the similarity metric. In this work, *K* = 5 is used for the majority voting since it achieved the best performance on validation data.
- **MusiteDeep**: MusiteDeep [8] is a recent method for kinase-specific phosphorylation sites prediction. The model uses a convolutional neural network with attention mechanism to extract features from the input peptide sequences without extensive human efforts for feature engineering. In comparison, we also ensured the MusiteDeep was trained on the same set of training data as our framework (i.e. data of training families and *k*-shot samples of the target family).

We did not compare our framework with other existing phosphorylation sites prediction methods for two reasons: i) The purpose of our experiments is to compare the prediction performance when only few-shot samples of the test family are given. Under this setting, we need to re-train the existing methods without giving them a large amount of data from the test family for model training. Most existing methods, including NetPhorest, GPS, NetPhos and PhosphoPredict, are available as a web server or released as pre-trained models, which probably have been trained with sufficient training samples. Therefore, it is difficult to evaluate the prediction performance of these methods in a fair setting. ii) For existing methods we are comparing to, MusiteDeep is open-sourced with codes allowing us to train customized models. In addition, recent works [14, 16] showed that MusiteDeep is one of the state-of-the-art methods, and we think it is a representative method of existing prediction approaches for phosphorylation sites prediction.

### 5.3 Few-shot learning performance

The first experiment is to evaluate our framework’s ability to use the few-shot samples to improve its prediction performance. After training the base predictor with the training families in the meta-learning phase, we directly evaluate the phosphorylation sites prediction performance for the base prediction on the *target* kinase family. We then further fine-tuned the base predictor, in the few-shot learning stage, using 10-shot samples from the *target* kinase family. We compared the prediction performances of the predictor prior to and after the few-shot learning phase (Figure 3). We found that with the broadly applicable model representation learned through the meta-learning phase, the predictor can rapidly adapt to a new family and capture its specificity. We observed the prediction performance of our framework was quickly improved after training on the 10-shot samples of the target family. This result suggests the potential of our framework in capturing the specificity of a novel kinase family, even if there are very few phosphorylation sites known for this family. We also note that the improvements do not solely come from the 10-shot sample but also the general representation learned by our framework in the meta-learning phase. We show in Supplementary Fig. S2 that only training the model with the *k*-shot samples was not enough to capture the specificity of a target family, emphasizing the importance of the meta-learning phase to achieve transferability.

**Figure 3:**
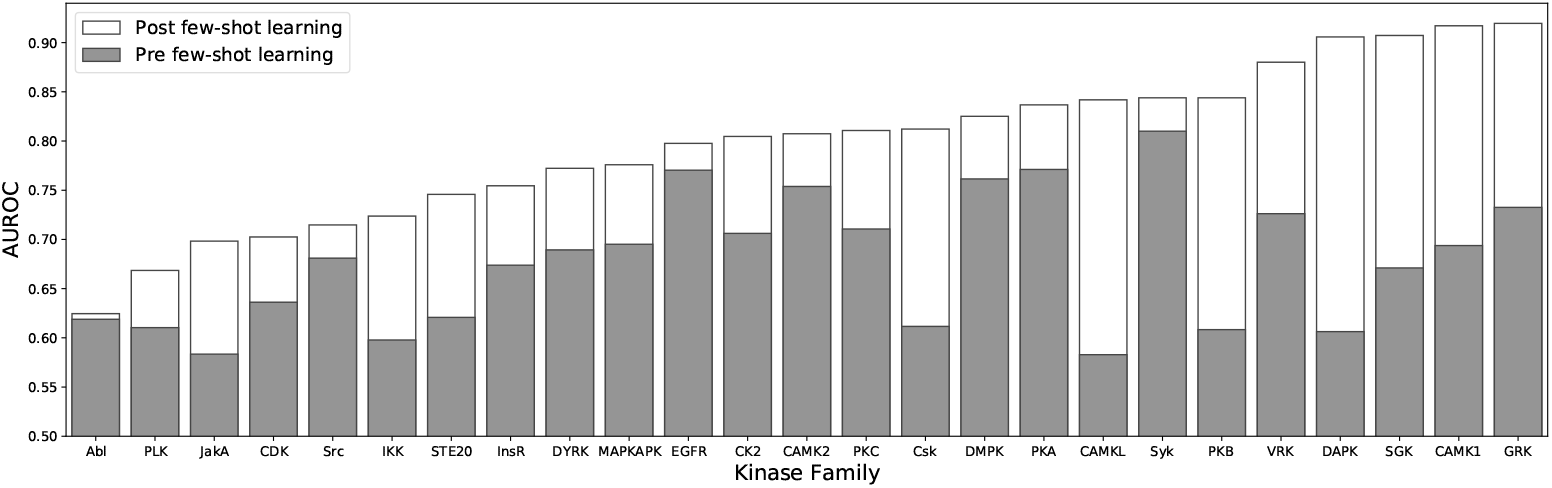
Improvements of prediction performance using few-shot learning. Pre few-shot learning: prediction performance of the model that was meta-trained on all training families but prior to the few-shot learning using the few-shot samples of the test family. Post few-shot learning: prediction performance of the model that was meta-trained on all training families and fine-tuned using the 10-shot samples of the test family in the few-shot learning phase.

### 5.4 Comparisons with baseline methods

Next, we compared our framework with the baseline methods. We varied the value of *k*-shot from 0 to 10 (0-shot means the model was trained on training family only), and for each value of *k*, we randomly sampled *k* samples from the target family and used the remaining samples as test data. The process was repeated for 50 times for each value of *k*. We used the AUROC and AUPRC scores as the evaluation metrics and showed the results in Fig. 4. We first observed that MetaKinase outperformed other methods for each value of *k* in terms of both AUROC and AUPRC scores. In addition, while other methods had relatively similar prediction performance as the number of *k*-shot increased, we observed that the improvement was clear for MetaKinase when more *k*-shot samples were provided. Our framework also achieved fast adaption to a target family. For example, the predictor had a 0.316 AUPRC score when using 2-shot samples in the few-shot learning phase, which was closed to AUPRC score achieved with 10-shot (0.338). These results demonstrated the transferability and fast-learning ability of MetaKinase.

**Figure 4:**
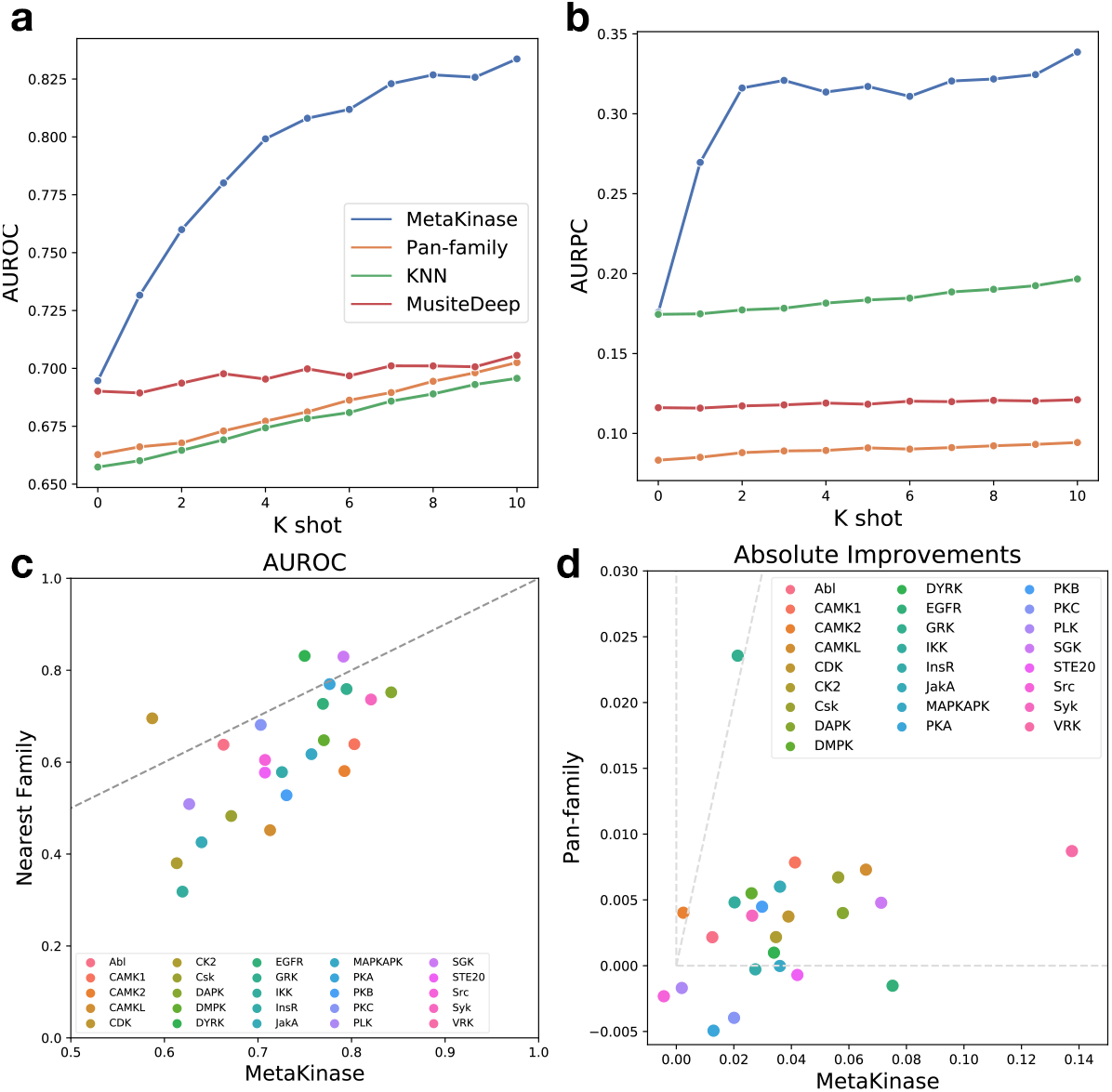
(a)-(b) Evaluation of few-shot learning. MetaKinase was trained with data of multiple kinase families in the meta-learning phase and fine-tuned in the few-shot learning phase using *k* samples of the test family for *k* = 1, 2, …, 10. For supervised-learning methods Pan-family, KNN, and MusiteDeep, the models were trained using data of training families and the *k*-shot data of the test family. The AUROC and AUPRC scores were used as the evaluation metric. Prediction performances were averaged over all 25 kinase families and 50 random samples of *k* shot data. See Section 5.1 for details of experiment settings. (c) Generalizability comparison between our method (0-shot) and the model trained on the nearest family of the target family (in terms of BLOSUM62 sequence similarity). (d) Effectiveness comparison between our method and pan-family method in using the first shot of the target family.

To further analyze the transferability of MetaKinase, we evaluated its prediction performance with 0-shot sample. This directly assesses the prediction power of the learned representation from the meta-learning phase. Here, we compared our framework with the nearest-family approach, where the model was trained on the most similar family of the target family. For each test target family, we computed all pairwise BLOSUM62 sequence similarities of any two kinases across the two families and took the family with the maximum average similarity as the nearest family of the target family. We used the same architecture as in MetaKinase for the nearest family approach. Comparison results for all families are shown in Fig. 4c. Even without training on *k*-shot samples, the predictor trained with data of training family already had a good prediction performance on the target family, suggesting that the model representation learned via the meta-learning phase is broadly applicable to each family and can achieve relatively good prediction performance. In addition, MetaKinase achieved higher AUROC scores than the nearest family approach and we conclude that integrating information from more families, instead of using the nearest one, is helpful to find a model representation with better transferability.

We also evaluated the effectiveness of MetaKinase in making use of the *k*-shot samples. To this end, we computed the absolute AUROC improvements achieved by MetaKinase and the pan-family approach for using 1-shot sample over using 0-shot (Fig. 4d). We found that our framework was sensitive to the first-shot samples seen in the few-shot learning stage and the prediction performance was clearly improved. The improvements achieved by our framework was significantly higher than that of pan-family approach (one-sided P-value 4.6 × 10^−7^4), which suggested our framework is sample-efficient in the few-shot learning stage.

### 5.5 Fast specificity learning by MetaKinase

As an exploratory analysis, we visualize the specificity captured by MetaKinase when using different numbers of few-shot samples. Here we used the specificity characterization of the InsR kinase family as an example. We first meta-trained the model using data of all families excluding InsR, and then fine-tuned the model in the few-shot learning stage, using 0-shot, 1-shot, 5-shot, and 10-shot, respectively. After the model was fine-tuned using a specific number of few-shot samples, we applied it to predict on the test dataset and extract the top 5% highest scoring peptides to visualize the logo representation. We showed in Fig. S3 the specificities captured by MetaKinase after trained on 0-, 1-, 5- and 10-shot samples, as well as the specificity derived from experimentally verified peptides. We observed that the specificity captured by MetaKinase gradually approximate the true specificity as the number of few-shot *k* increases. At 0-shot, the predictor was only trained on data samples of other families and the specificity captured (Fig. S3a) was clearly different from the true specificity (Fig. S3e), presumably because what the model tried to capture is a general specificity for all training families but less specific to the target family. After the model had been trained with few-shot samples of the target family, we found that the model was able to capture the specificity for the target family. For example, position 3 at 1-shot, positions 1 and 3 at 5-shot, and positions −7, −2 and 3 at 10-shot. These visualizations illustrate that our framework is able to rapidly capture the specificity of a target family using only a small number of data samples.

## 6 Discussion

In this paper, we introduced a generic framework for protein-peptide binding prediction and implemented MetaKinase for kinase specificity prediction as a proof-of-concept. We demonstrate its applicability in predicting the phosphorylation sites of protein kinases. We utilized meta-learning and few-shot learning strategy to mitigate the data scarcity issue in characterizing the specificity of less-studied kinases. The framework can take an arbitrary gradient-based model as the base predictor. MetaKinase is capable of learning a general representation for each kinase family and quickly capture the specificity of a target family using a few data samples, without overfitting. We demonstrated that MetaKinase outperformed existing approaches in predicting kinase-specific phosphorylation sites when training samples are limited. The framework is sample-efficient and can quickly capture the specificity of a target family using a few data samples. There are several directions for future study. For example, our framework, in principle, can be applied to predict other types of protein-peptide binding, including the phosphorylation sites of SH2 domain [28], the binding of MHC to neoantigens [29].

## Acknowledgements

This work was supported in part by the NSF CAREER Award, the Sloan Research Fellowship, the PhRMA Foundation Award in Informatics, and the CompGen Fellowship.

## Supplementary Information

**Figure S1:**
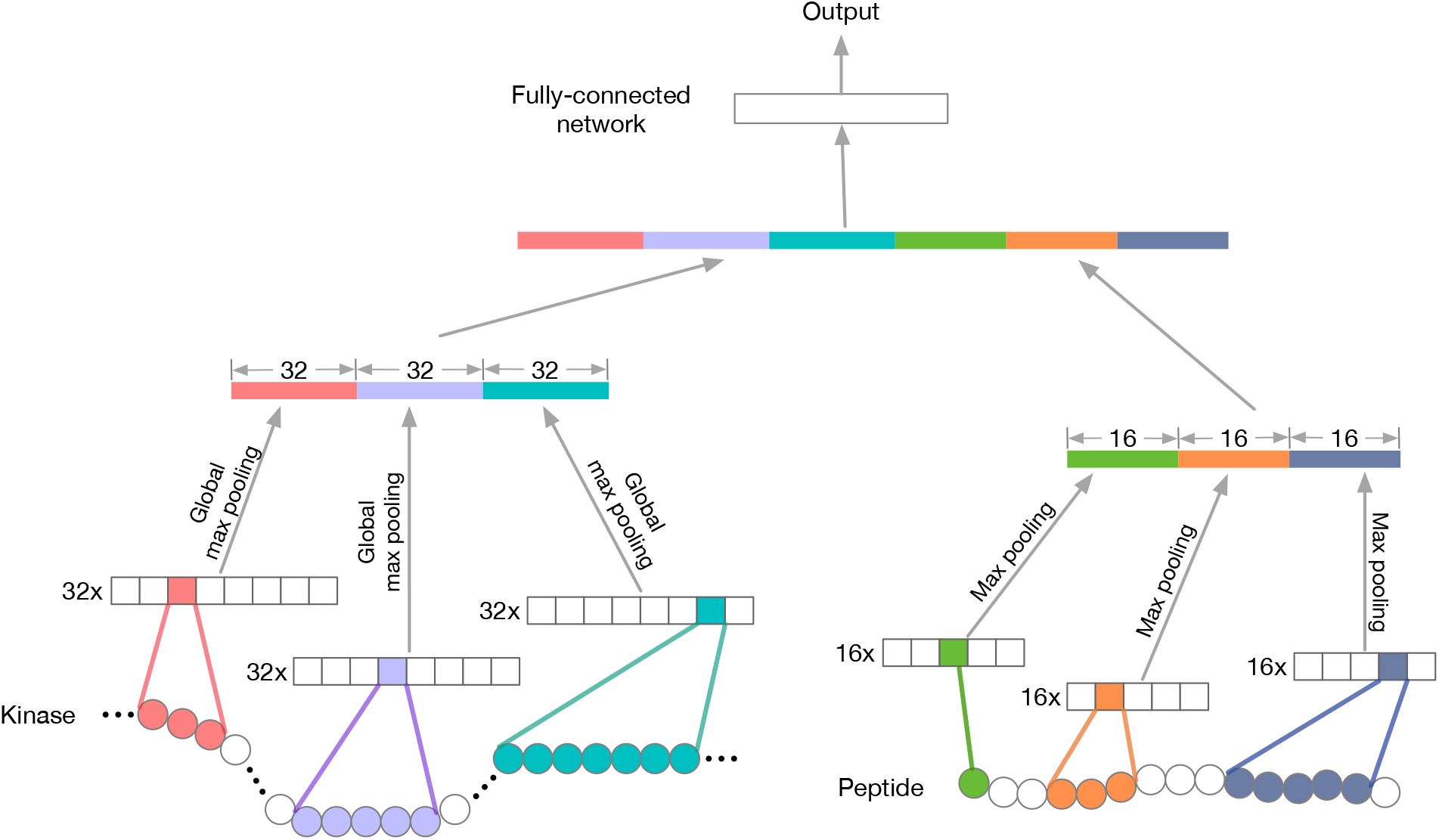
Model architecture of base predictor. We utilize a dual-channel deep neural network as the based predictor. The input of the left channel is the kinase domain sequence (~ 250 amino acids) while the peptide sequence (15 amino acids) is fed into the right channel. We use BLOSUM62 matrix [27] to encode the input sequences. Each channel consists of three convolution neural networks (CNNs) with different kernel sizes. 32 filters of size 3, 5 and 7 make up the left channel. Meanwhile, 16 filters of length 1, 3 and 5 compose the right part. Then, a global pooling operation is performed to select the maximum activation value in each of 32 × 3 = 96 filters in the left and 32 × 2 = 64 in the right. After concatenating the outputs from both channels, a fully-connected layer produces the final prediction.

**Figure S2:**
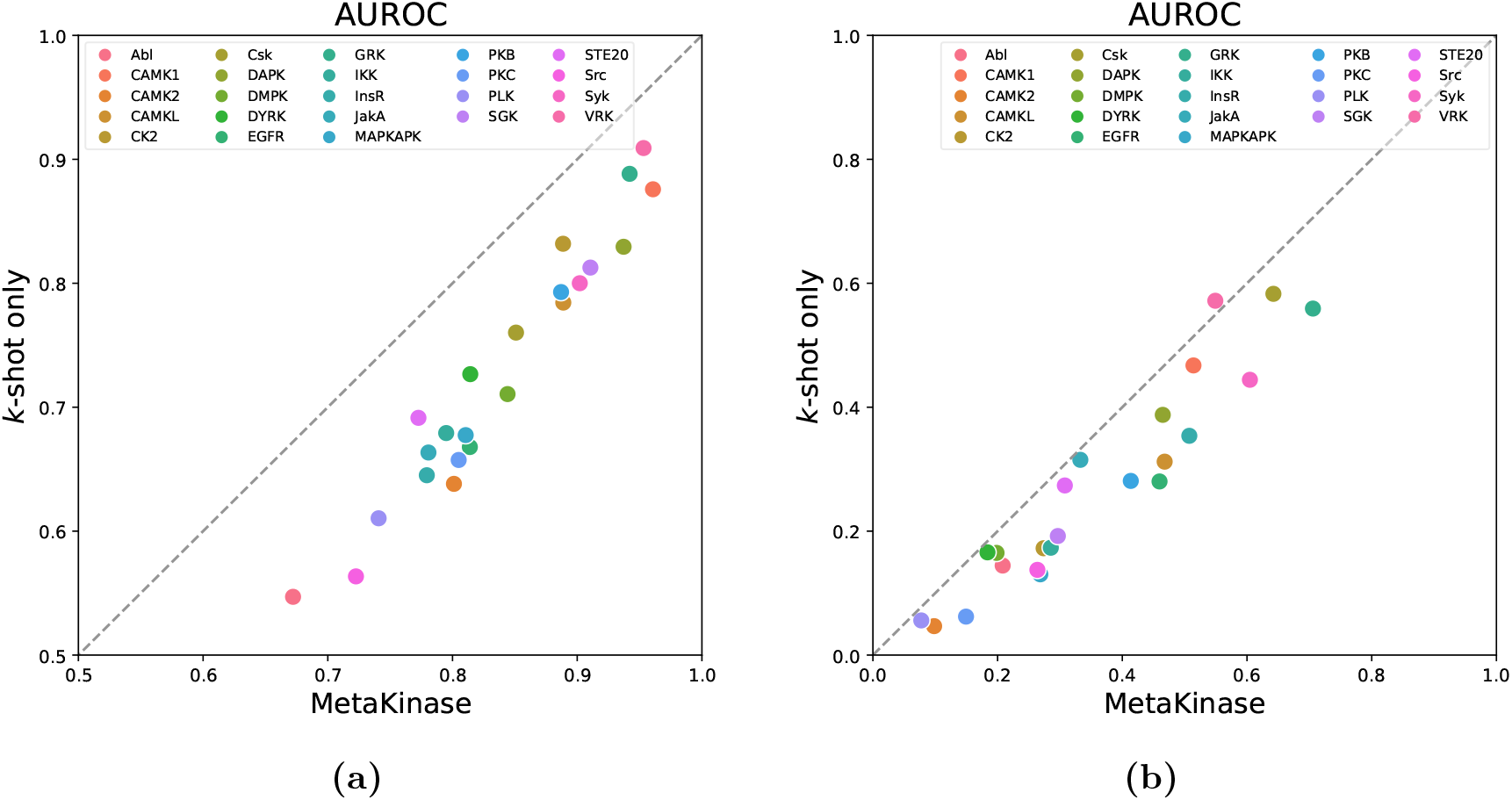
Comparison between MetaKinase and the model trained with *k*-shot samples. MetaKinase was trained on the *N* − 1 training families through a meta-learning phase and then fine-tuned using 10-shot samples of the target family in the few-shot learning stage. The model trained with 10-shot samples was trained with only the 10-shot samples using in a supervised-training manner.

**Figure S3:**
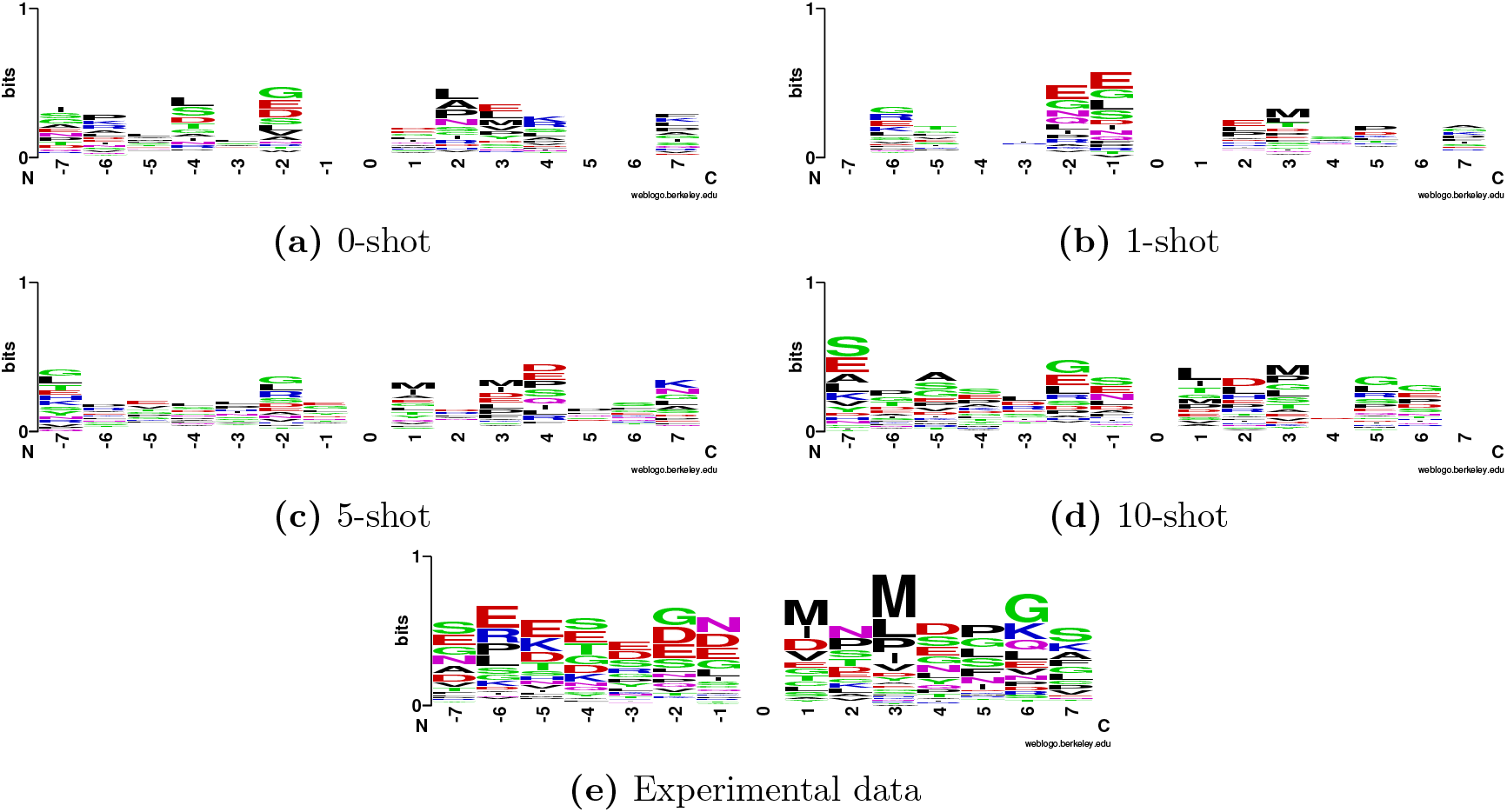
Sequence logo representation of specificities of InsR kinase family. (a)-(d) Specificities captured by models fine-tuned with 0, 1, 5 and 10 samples, respectively. (e) Specificity derived from experimentally verified phosphorylation sites. The phosphorylation residue (position 0) of InsR family is always Tyrosine (Y) and is omitted in the logo representation.

